# THINGS: A database of 1,854 object concepts and more than 26,000 naturalistic object images

**DOI:** 10.1101/545954

**Authors:** Martin N. Hebart, Adam H. Dickter, Alexis Kidder, Wan Y. Kwok, Anna Corriveau, Caitlin Van Wicklin, Chris I. Baker

**Author notes:** Correspondence should be addressed to: Martin Hebart. **Author contribution** M.N.H. and C.I.B. designed the research, M.N.H., A.H.D., A.K., W.Y.K., and C.W. identified and selected object concepts, M.N.H., A.H.D., A.K., W.Y.K., A.C. and C.W. identified, selected and post-processed object images, M.N.H. and A.H.D. carried out analyses, all authors wrote the paper or contributed significantly to its revision. **All data available at:** http://doi.org/10.17605/osf.io/jum2f.

## Abstract

In recent years, the use of a large number of object concepts and naturalistic object images has been growing strongly in cognitive neuroscience research. Classical databases of object concepts are based mostly on a manually curated set of concepts. Further, databases of naturalistic object images typically consist of single images of objects cropped from their background, or a large number of naturalistic images of varying quality, requiring elaborate manual image curation. Here we provide a set of 1,854 diverse object concepts sampled systematically from concrete picturable and nameable nouns in the American English language. Using these object concepts, we conducted a large-scale web image search to compile a database of 26,107 high-quality naturalistic images of those objects, with 12 or more object images per concept and all images cropped to square size. Using crowdsourcing, we provide higher-level category membership for the 27 most common categories and validate them by relating them to representations in a semantic embedding derived from large text corpora. Finally, by feeding images through a deep convolutional neural network, we demonstrate that they exhibit high selectivity for different object concepts, while at the same time preserving variability of different object images within each concept. Together, the THINGS database provides a rich resource of object concepts and object images and offers a tool for both systematic and large-scale naturalistic research in the fields of psychology, neuroscience, and computer science.

## Introduction

Two central goals in cognitive neuroscience are to elucidate how we recognize the objects in the world around us and how we are able to form categories based on our percepts and our semantic knowledge. Reaching these goals requires us to overcome two challenges. First, we need to understand how the visual system identifies the relevant object features from the almost infinite number of possible object appearances. Second, we need to understand how observers use this information about the object and integrate it with their object knowledge to uniquely identify its semantic content while distinguishing it from the thousands of other concepts to which it could belong. Computer science, in particular computer vision, is grappling with similar challenges for artificial visual systems. Crucially, for any study involving object concepts or the visual presentation of object images, both the selection of concepts and the images depicting those concepts can strongly influence the results of a study and the conclusions that are drawn.

### The need for large-scale systematic sampling of object concepts and naturalistic object images

In recent years, in the fields of psychology, neuroscience and computer science there has been a growing interest in utilizing a wide range of object concepts and naturalistic images (Deng et al., 2009; Einhäuser & König, 2010; Felsen & Dan, 2005; Oliva & Torralba, 2007; Pereira et al., 2018). This interest arises primarily from the goals of (1) achieving an ecologically valid understanding of visual and semantic cognition, and (2) building generalizable computational models of object recognition (e.g. Krizhevsky, Sutskever, & Hinton, 2012) and semantic knowledge (e.g. Mikolov, Yih, & Zweig, 2013; Pennington, Socher, & Manning, 2014). Thus, the availability of a wide range of systematically sampled object concepts and naturalistic object images promises not only to benefit large-scale experimental and computational approaches to the study of visual and semantic cognition; it also offers classical small-scale experimental approaches the possibility of selecting a more representative set of concepts and object images for testing specific hypotheses regarding cognition, behavior, and neural representations (Pereira et al., 2018).

Influential databases of object categories and concepts (Battig & Montague, 1969; Van Overschelde, Rawson, & Dunlosky, 2004) cover only a selective subset of the objects found in the everyday environment. In contrast, the lexical database WordNet (Fellbaum, 1998) offers a highly-systematic taxonomy of a vast range of words and their meaning, but its great detail makes it challenging to select a representative set of object concepts. Understandably, for many experiments the manual selection of individual objects or object categories is still common practice.

For object images, there are numerous databases from research in psychology and neuroscience, which most commonly contain line drawings of objects (Snodgrass & Vanderwart, 1980) or photographs of objects cropped from their natural background (for a review, see Brodeur, Dionne-Dostie, Montreuil, & Lepage, 2010; Brodeur, Guérard, & Bouras, 2014). However, naturalistic object context has been shown to play an important role in object recognition (Oliva & Torralba, 2007), and cropped object images may overemphasize the role of shape in neural representations of objects (Bracci, Daniels, & de Beeck, 2017; Bracci & de Beeck, 2016; Coggan, Liu, Baker, & Andrews, 2016; Proklova, Kaiser, & Peelen, 2017; Proklova, Kaiser, & Peelen, 2016). In addition, such databases usually offer only one example per object concept, while the use of multiple object examples is desirable for measuring generalizable representations. In the machine learning community, several large-scale object image databases have been developed and are commonly used, with up to thousands of image examples per concept (Deng et al., 2009; Everingham, Van Gool, Williams, Winn, & Zisserman, 2010; Griffin, Holub, & Perona, 2007; Krizhevsky & Hinton, 2009). However, most of the images contained in these databases are too small to be of practical use in psychology and neuroscience experiments, and images vary strongly in aspect ratio and quality and often still lack naturalistic backgrounds, making it challenging to use them (Chang, Pyles, Gupta, Tarr, & Aminoff, 2018). More recent databases (Kuznetsova et al., 2018; Lin et al., 2014; Zhou, Lapedriza, Khosla, Oliva, & Torralba, 2018) may offer higher quality, but are composed mostly of images containing multiple different objects at the same time or naturalistic scenes such as cities, beaches, or forests. In contrast to real-world scenes, many researchers are interested in using image databases focused on images of individual objects.

### Aim of the object concept and object image database

With this present work, we aim at providing researchers in the fields of psychology, neuroscience, and computer science with (1) a large-scale systematic sampling of object concepts, (2) a list of object categories derived from those concepts, and (3) a large set of high-quality color images of objects with naturalistic background. This database, together with similarity matrices based on semantic embeddings and activations in a deep convolutional neural network, will be made freely available for academic purposes in the final journal version of the paper. Until that time, a download link is available upon direct contact with the authors.

## Methods

### Data availability

The object concept and image database as well as all data and results are made publicly available through the Open Science foundation at http://doi.org/10.17605/osf.io/jum2f.

### Participants

We recruited a total of 1,395 workers from the online crowdsourcing platform Amazon Mechanical Turk for different tasks involved in the creation of this database, including object image naming and object categorization (see below). In the object naming task, workers were compensated $0.10 for 10 responses (in practice ~$7.50/h), and in the object categorization task $0.05 for 5 responses (in practice ~$5.00/h; all estimates based on the mode of the completion time). This research was approved by the NIH Office of Human Subjects Research Protections (OHSRP), and workers were compensated financially for their time.

### Identification of picturable and nameable object concepts

The selection and identification of the 1,854 object concepts used for this image dataset encompassed a three-step procedure that is described in more detail below. First, from an existing word database, we gathered a list of nouns that represent concrete, picturable object concepts. Second, we carried out word-sense disambiguation by assigning each noun one or several unique WordNet identifiers (“synsets”) that represent the meaning of this noun, thus allowing us to eliminate synonyms or identify nouns with multiple meanings. Third, we identified the subset of synsets that matches their use in everyday language, by selecting representative images for all synsets and testing how consistently they were named by human subjects. Note that the final list is not intended to be a complete and definite set of *all* picturable and nameable object concepts in the English language (see *Discussion*). However, the steps below constitute a systematic approach towards their selection and a detailed description of the different decisions involved. In addition, note that the actual number of selected nouns at each of these steps may deviate slightly from the numbers reported below, as in some cases during each selection step we identified items that were mistakenly kept but should have been excluded at earlier steps.

#### Step 1: List of concrete picturable nouns in American English

To compile a list of concrete picturable nouns, we used a list of American English lemma (Brysbaert, Warriner, & Kuperman, 2014) that contains concreteness ratings for ~40,000 words and two-word expressions (e.g. “ice cream”). For simplicity, we will collectively refer to these words or two-word expressions as words. In this list of American English lemma, concreteness ratings ranged from 1 to 5 and reflected the level through which the word could be experienced through one of the five basic senses (5: concrete, 1: abstract). The concreteness rating of each word is based on the average of 25 to 30 unique behavioral responses.

To select candidate lemma from this list, we applied two selection criteria. First, we restricted our selection to words that were tagged as nouns. We used part-of-speech tags provided with the concreteness ratings (Brysbaert, New, & Keuleers, 2012) and extended them using the British Lexicon Project (Keuleers, Lacey, Rastle, & Brysbaert, 2012), which offers multiple tags per word (e.g. “cook” both as noun and as verb). Second, we restricted our selection of words to those with a concreteness rating ≥ 4. This choice was based on a preliminary manual screening of the list, since we only rarely identified candidate words with a lower concreteness rating. Based on these criteria, we identified 8,671 nouns.

Following this initial selection, we manually screened the list based on a number of exclusion criteria (a list of the excluded lemma is provided with this database). The exclusion criteria are: (1) not a noun despite part-of-speech tag, (2) plural form when singular form with the identical meaning is found in the list (e.g. exclude “cats” when “cat” is present, but keep “glasses” when “glass” is present), (3) clear synonym (e.g. “motorcar” vs. “car”), (4) nouns that were clearly not nameable or too general (e.g. “equipment”), (5) navigable places (e.g. “garden”) including integral parts of buildings (e.g. “steeple”), (6) nouns referring to persons in certain roles (e.g. “doctor,” “pilot,” “audience”), with certain ethnic or cultural origin (e.g. “Indian”), or with certain color of skin (e.g. “albino”), (7) fictitious or extinct beings for which no real-world photograph can exist (e.g. “dragon,” “werewolf,” “dinosaur”), (8) relationship statuses (e.g. “father,” “granddaughter”) (9) body parts that are either internal organs or muscles (e.g. “liver”) or where the noun describes these as being parts of other body parts (e.g. “fingernail,” “forearm”), (10) bodily fluids (e.g. “urine”), (11) body parts of animals that do not serve as tools for humans (e.g. exclude “beak” but include “feather”), (12) non-visual but sensory nouns (e.g. “click,” “music”), (13) action nouns (e.g. “smack”), (14) times of day (“night”), (15) units and geometric figures (e.g. “quart,” “hexagon”), (16) fluids or light that cannot be ascribed to an object (e.g. “floodwater,” “moonlight”) or specific drinks that cannot be identified without a label (e.g. “whisky,” “gin”), (17) celestial bodies (e.g. “moon”), (18) specific drugs, pharmaceuticals and chemical compounds, (19) parts of objects that are difficult to describe in isolation (e.g. “seam,” “rim”), (20) fabrics and unspecific surface materials (e.g. “tweed,” “teakwood”), (20) brand names unless objects of other brands share this name (e.g. exclude “iPad” but include “jeep”) and (21) nouns denoting or implying sexual content or nudity. In addition to these exclusion criteria, post-hoc we chose to exclude all nouns that require text for identification of the object (e.g. “lexicon”), or that refer to objects that necessarily contain text (e.g. “newspaper”), since for an image database the identification should not rely on text or be biased by the written text. Finally, we excluded a small set of nouns that were not in WordNet (see below) and that we judged as very unusual (e.g. “wishbone”). This selection left us with a list of 3,397 words.

#### Step 2: Word sense disambiguation through assignment to WordNet synsets

Each of the nouns identified in the previous step may carry more than one meaning, making it ambiguous as to which concept the noun represents (e.g. “bat” as an animal or “bat” as a sports item). In addition, the list may contain synonyms, i.e. different words with the same meaning that could be merged (e.g. “couch” and “sofa”). To identify all unique picturable meanings from this list of words, we used WordNet to carry out word-sense disambiguation, i.e. the assignment of words to unique senses. In WordNet (Fellbaum, 1998), a meaning or sense is referred to as a synset, which comes with a unique identifier (synset ID), a list of synonyms, and a definition. Note that while the coverage of word meanings in WordNet is extensive, some meanings are not covered, and others are represented by multiple synsets that could in principle be merged.

To identify unique synsets, we first compiled all candidate synsets for each word in our list. For words with only one synset, the assignment between word and synset did not require any disambiguation. For those words, we removed synsets whose definition did not represent a picturable object and that had mistakenly not been eliminated in the previous step. For all remaining words, we created a graphical user interface (GUI) to present each word alongside all of its synsets (Fig 1). In the GUI, the reference word appeared on top, alongside all candidate synsets with different synonyms, definitions and, when available, three matching images extracted from ImageNet (Deng et al., 2009). For every word, two independent raters selected the meaning(s) that best matched the word. Two raters carried out this assignment for one half of the words (even numbers), and another two raters for the other half (odd numbers). The mean inter-rater agreement was 82.53%, and disagreements were resolved by a third rater. Based on these ratings, we merged synonymous words by picking the synonym used most frequently, using the word frequency provided in the Corpus of Contemporary American English (Davies, 2008). In addition, we assigned words with multiple meanings separate identifiers (e.g. “bat1” and “bat2”). This process left us with a total of 3,228 object concepts based on the meaning of different words.

**Fig 1.**
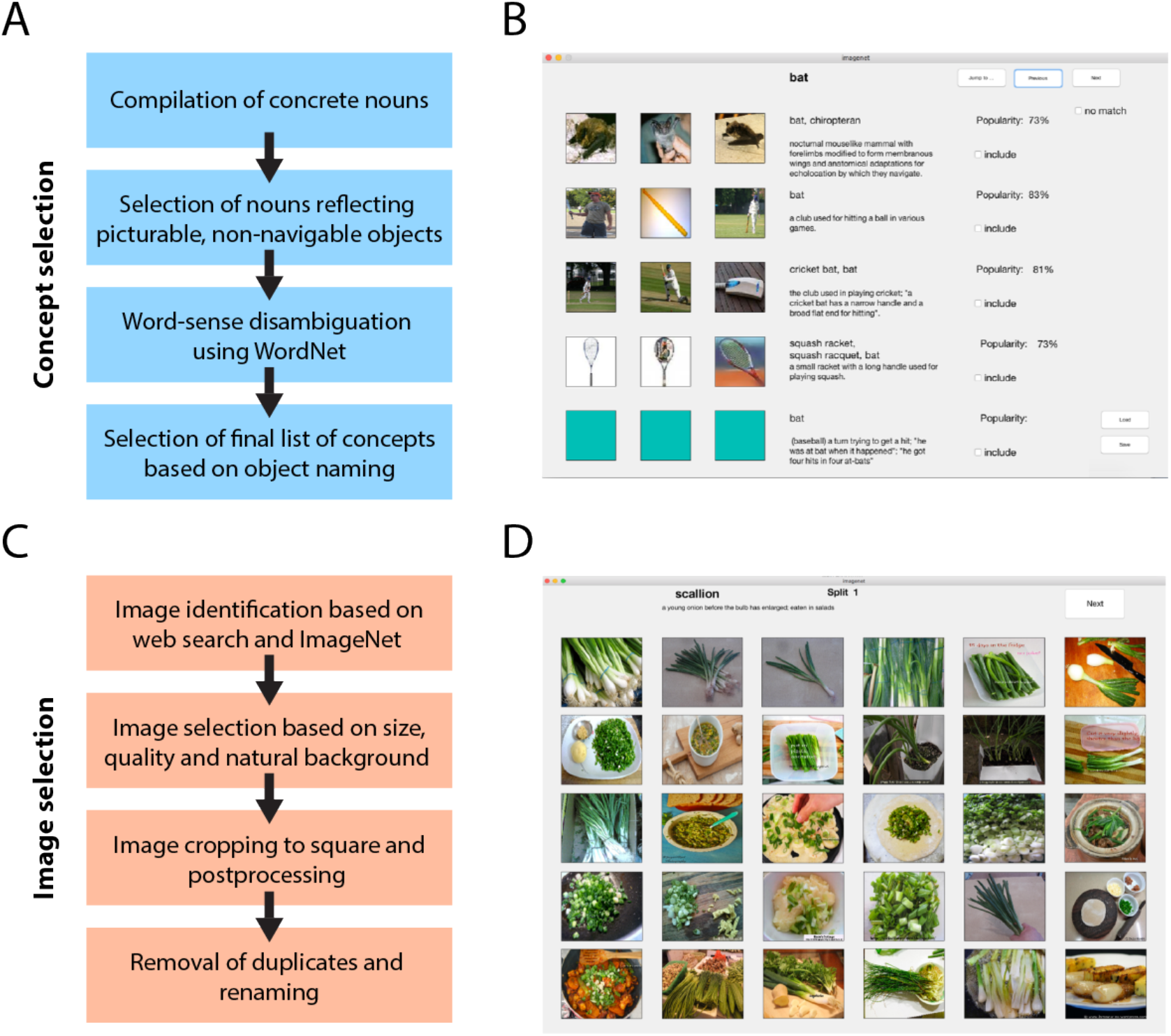
Concept and image selection procedure and graphical user interfaces (GUIs) used for concept and image selection. **(A)** Procedure for selection of 1,854 concepts in the THINGS database. **(B)** GUI used for word-sense disambiguation based on WordNet senses (synsets) and example images from ImageNet. **(C)** Procedure for selection of 26,107 object images in the THINGS database. **(D)** GUI used for initial manual selection of candidate images with sufficient quality.

#### Step 3: Object naming task to identify picturable object concepts

Using the list of concepts from the previous step, we identified representative images depicting those concepts and conducted an object naming task. The aim of this task was to identify the set of objects for which participants use the intended object concept and separate them from those for which they use a different one. The reasoning behind this approach is twofold. First, including object concepts according to how they are named improves their comparability and prevents ambiguity with respect to object knowledge. Second, for a database of object images, an observer should be able to both recognize an object and identify the object concept. There are several reasons why there might be disagreement between the concept and the name an observer uses for the object in the image. First, some of these words are quite specific and reflect a subordinate category level (e.g. “blue jay”) which may not correspond to the description commonly used by human observers (e.g. “bird”). Indeed, this procedure has been used previously, and it was demonstrated that naming results commonly reflect the basic level in a naming task (Rosch et al., 1976). However, we did not restrict the selection of concepts to the basic level, but specifically chose concepts based on their everyday use in language. Second, some of the concepts represent object parts (e.g. “hat ring,” “lip”) which can only be shown in the context of an object when zooming in, but which may lead observers to nevertheless focus on the object (e.g. “hat,” “mouth”). Finally, naming might be highly inconsistent or incorrect, indicating that observers have difficulties identifying the object concept or disagree which concept to use.

We tested object naming by selecting a representative object image with natural background for each of the 3,228 object concepts and presenting them to workers on Amazon Mechanical Turk (*n* = 445, mean number of responses per worker: 86.29). Each image was shown to 10 workers who were asked to label the object or “thing” that was most prominent in the image by typing it into a box below the image. An additional set of 10 responses was collected for ambiguous cases. The answers were corrected for spelling errors and plural forms, and different synonyms of a synset were labeled as reflecting the same concept. Our inclusion criteria were rather liberal, i.e. we opted to rather include a concept than to falsely exclude it. All responses are made publicly available with the database, including links to the representative images. Exclusion criteria for object concepts were: (1) if the expected label was provided only once, (2) if the expected label was provided only twice while a different label was consistently provided at least five times, (3) if the expected label was provided only three times while a different label was consistently provided at least six times. Criteria for collecting an additional 10 responses were: (1) If the expected label was provided twice while a different label was consistently provided less than five times, or (2) if there were an equal number of responses with the expected label and a different label. In addition, we manually inspected all concepts marked for exclusion, and when there was any doubt that the image was not representative enough, we selected a new image and retested the concept. After collecting the additional responses, all concepts that were still not clearly included or excluded were left in. Based on this approach, an additional 1,374 concepts were excluded. Of those, 706 were named at a higher taxonomic level, 630 were named inconsistently, and 38 referred to a whole object when the object concept reflected a part. This left us with a final number of 1,854 object concepts.

### Identification of higher-level object categories

Having identified the final set of 1,854 object concepts, we sought to identify a set of higher-level object categories. To this end, we followed two strategies. In line with the approach of identifying the concepts based on their use in everyday language when naming concrete objects, we chose a similar “bottom-up” strategy of having workers on Amazon Mechanical Turk identify the categories each object belongs to. However, this approach likely results in errors based on incorrect beliefs (e.g. that a peanut is a nut rather than a legume) and may lead to incomplete categories. Hence, we complemented this strategy with a “top-down” approach, identifying members of the dominant categories based on the taxonomy of word senses inherent in WordNet.

#### Bottom-up determination of object categories

For the bottom-up strategy, we conducted a two-step procedure on Amazon Mechanical Turk, first asking workers to propose candidate categories for those objects and second asking a separate group of workers to select the most appropriate category from those candidates. In more detail, in the first step, for each concept and representative image we asked 20 workers to provide the category this object belongs to (*n* = 427, mean number of responses per worker: 43.41). The instructions included clear examples using abstract concepts and categories, in order not to set a strong baseline for what results are expected for concrete objects (e.g. “blue” is a “color,” “Susan” is a “name” or “female name”). We excluded responses that simply repeated the original concept, but kept all other responses, including arbitrary categories (e.g. “sports” rather than “sports item” or “sports equipment”). In addition, we corrected all responses for spelling errors, and converted plural form to singular form (unless the original concept was a plural noun, such as “glasses”).

In the second step, to identify representative categories from those resulting in the first round, we asked another 20 workers per concept (total *n* = 523) to provide a category for the object. This time, however, in addition to the word representing the concept and the representative image, we showed workers the answers that had been provided by workers in the previous step, in random order. Workers were instructed that some of the responses were incorrect, that they could combine previous responses, and that they could use their own category if they deemed all responses inappropriate. Responses of one worker were removed who carried out an unusually large number of responses with unusually fast response times (mean number of responses per worker: 67.22, number of responses of excluded worker: 1,990). All remaining responses in this second step were again corrected for spelling errors, and plural form was converted to singular form. In addition, we unified responses that were synonymous (e.g. “beverage” and “drink”) and removed qualitative statements (e.g. “large,” “small,” “hot,” “cold,” etc.), colors, and place of origin (e.g. “Asian,” “French”). Responses were automatically assigned to a higher-level category if (a) there were 11 or more consistent responses or (b) there were at least 5 consistent responses while less than half that number were a consistent alternative. All assignments were later checked manually for accuracy, and higher-level categories were assigned to all concepts based on a summary of the provided categories or the use of synonymous categories. Finally, from the list of all categories, we identified categories that were used synonymously or overlapped strongly (e.g. “kitchen tool” vs. “kitchen utensil”) and merged them.

#### Top-down determination of object categories

For the top-down strategy, we used the taxonomy in WordNet to identify all concepts that were subordinate to the 27 high-level categories identified in the previous step. To this end, we first identified the synsets of the high-level categories. Of those categories, 23 were found in WordNet, and we used the category “decoration” in place of “home décor,” due to the strong overlap between the members of both categories, making it a total of 24 categories. For each of the 1,854 object concepts, we recursively ascended each branch of the WordNet tree until we reached the top synset. We then assigned the object concept to any of the 24 categories that was crossed while ascending through WordNet. Note that the category “car part” does not contain any subordinate entries in WordNet, i.e. in the *Results* section, the number of categories is described as 23.

### Selection of images for object image database

For the final object image database, the goal was to identify at least 12 images for each of the 1,854 object concepts, i.e. a minimum of 22,248 images. Image selection and postprocessing (e.g. cropping) were conducted by the authors. Note that below, the identification of candidate images and postprocessing are described as sequential steps. However, the process of selecting object images went through multiple cycles until a sufficient number of suitable images had been identified for all concepts.

#### Image selection criteria

The goal for this object image database was to select high-quality, naturalistic images of objects belonging to the 1,854 object concepts. Each image was a photograph of one or several examples of a given object concept that was cropped to square size. Since the selection criteria (detailed below) required us to exclude the majority of candidate images, for some concepts it was difficult to find examples of suitable quality. For those concepts, we decided to slightly loosen some of these selection criteria. In the following, when we write of “exclusion,” this refers to strict exclusion criteria for images, while the term “avoiding” refers to exclusion criteria that were less strict, depending on how difficult it was to find object examples.

The objects were generally chosen to be the central component of the image, and while additional objects of other concepts were allowed to be present in an image, they were not allowed to dominate the image. For example, the presence of human body parts was permitted in images of clothing; however, we avoided images showing human faces due to their strong salience (with the exception of concepts defined by human faces, e.g. “face,” “man,” “woman”). Since each image was cropped to a square in order to standardize the image size, we focused on identifying images that would still show the majority of an object after cropping.

The images were photographs with a minimum size of 480 × 480 pixels, but most were 600 × 600 pixels or larger (see Fig 3B). We selected images of objects with naturalistic background, i.e. we avoided images with uniform background and excluded images for which the natural background had been removed or which had been modified in a non-naturalistic and recognizable fashion. We avoided images in which the object was blurry or for which lighting was over- or underexposed, and we excluded images with non-naturalistic colors (including grayscale), strong color filters, watermarks, borders, or other text added to the images. Finally, we specifically avoided text that naturally appeared within the images (e.g. on a book), especially when the text allowed the identification of the object concept. However, for some object concepts, this was difficult or impossible to avoid (e.g. “police car”).

#### Identification of candidate images

Due to the large number of object concepts and images as well as the strict selection criteria, we pursued several strategies for identifying suitable images. First, we automatically selected 30 candidate images per concept from each of the image search engines Google Images, Bing Images and the photography website Flickr. Then, we manually selected candidate images, using a custom-made graphical user interface (GUI) written in MATLAB (Mathworks, Natick). Since this strategy did not yield a sufficient number of high-quality object images for most object concepts, we decided to directly identify and download candidate images through manual web searches, with a focus on images from Google Images, Bing Images, and the online-auctioning platform eBay. We accelerated the process by opening a set of candidate images in the browser and using a bulk image downloader that allowed us to select images with sufficient size (https://chrome.google.com/webstore/detail/fatkun-batch-download-ima/nnjjahlikiabnchcpehcpkdeckfgnohf). Search terms generally focused on the word reflecting the object concept, in some cases using a translation into Spanish, German, Russian, Chinese, or French. In addition, we identified a set of candidate images of sufficient size from ImageNet (Deng et al., 2009), which had been prescreened and selected for the object image database “ecoset” (Mehrer, Kietzmann, & Kriegeskorte, 2017), followed by manual selection with the image selection GUI. Finally, several images taken by the authors were added to the database.

#### Image cropping, removal of images with border, and manual quality check

Following the manual selection of suitable candidate images, we cropped objects to square size, using a separate custom-made GUI (written in MATLAB) to accelerate the process (Fig 1). This GUI allowed us to select relevant image parts for sufficiently large images, crop images to square size, and exclude bad candidate images. Further, the GUI was written to prevent the selection of image parts that were smaller than 480 × 480 pixels. Following this image cropping, we identified and corrected images that still contained a small uniform border by searching for uniform pixel intensities at the edge of the image. Finally, we manually screened all cropped images a second time to identify and remove images with low quality or those matching other exclusion criteria.

#### Semi-automatic identification of highly similar or duplicate images

A small number of candidate images within a concept were duplicates, photographs of the same object from a slightly different perspective, or images with different object examples but identical background. To identify highly-similar images or duplicates, we passed all images through the deep convolutional neural network VGG-16 (Simonyan & Zisserman, 2014), which had been pretrained on 1,000 ImageNet concepts as implemented in the toolbox MatConvNet (http://www.vlfeat.org/matconvnet/pretrained/). We reasoned that highly similar images would produce similar activation vectors at different levels of generalization.

For the identification of those images using VGG-16, we focused on the five pooling layers and two fully-connected layers, i.e. a total of seven layers.

In the following, we describe the process for the first layer, which we repeated separately for all seven layers. For all object images, we computed the activations for the current layer and vectorized them. Subsequently, within each of the 1,854 object concepts, we calculated the Pearson correlation coefficients between all pairs of vectors. For example, for an object concept containing 12 object images, this step yielded 66 correlation coefficients. Finally, we used MATLAB to display the object pairs subsequently, starting with the pair with the largest correlation coefficient and subsequently moving down correlation coefficient size. We manually checked the first 1,500 pairs and removed images that were duplicates, were the same object taken from different angles, or had an identical background. Screening the first 1,500 pairs proved to be effective, and not a single image was removed within the last 100 of those image pairs. We repeated this process for all seven layers. Since different network layers produced similar duplicate candidates, we excluded pairs that had been screen previously, but still screened an additional 1,500 pairs manually per layer.

#### Assignment of image order and image name in final database

As the final step, we converted all images to jpeg-format, determined the final order of object images, and standardized their filenames. As described above, the goal for the final database was to have at least 12 images per concept. For the reason of simplicity, we deemed the first 12 images of each concept to be the most relevant ones. The first image of each concept was the reference image used in the object naming task, unless it was not of sufficient quality and had been excluded (number of excluded reference images: 191). All other images were initially sorted by image size. Then, we shuffled the order of the 11 largest images. For all concepts with more than 12 images, we separately shuffled the order of the remaining images. Since for comparison to computational vision algorithms it is often important to know which images were used for training an algorithm and since ImageNet is commonly used, all images chosen from ImageNet were labeled with the letter n. All reference images were labeled with the letter b, and all other images with the letter s.

### Similarity matrices from computational models of semantics and vision

#### Semantic embedding based on synset vectors

Semantic embeddings provide a low dimensional vector representation of words based on their co-occurrence statistics in large text corpora that approximate the relationship between word meanings. Recent developments in word embeddings based on shallow neural networks, in particular word2vec (Mikolov et al., 2013), have led to strong improvements in performance. A recent modeling approach (Pilehvar & Collier, 2016) combined word2vec with knowledge about different senses based on WordNet synsets, providing separate vectors for different meanings of words. Here, we extracted those synset vectors (1) to create a similarity matrix in order to visualize the semantic distribution of different concepts, and (2) to provide a quantitative basis for the selection of a representative subset of concepts based on their semantic similarity and (3) to offer a resource for researchers who intend to use them alongside the concepts. In short, for all synsets, we extracted 300-dimensional synset vectors from those provided by Pilehvar and Collier (2016) (https://pilehvar.github.io/deconf/) that had been trained with word2vec on the Google News Corpus (https://code.google.com/archive/p/word2vec/). For words not represented in WordNet or missing in the synset vector representations, we chose the original word2vec model, but rescaled word vectors to have the same standard deviation as synset vectors since synset vectors were reduced in variance. Finally, we calculated a similarity matrix between words using the Pearson correlation (the common cosine distance and Euclidean distance led to very similar results, all *r* > 0.96).

#### Deep convolutional neural network activations for all object images

Deep convolutional neural networks (CNNs) offer a computational model of object recognition and – due to their excellent performance – have become very popular models not only in the field of computer science, but also in psychology and neuroscience (Cichy, Khosla, Pantazis, Torralba, & Oliva, 2016; Khaligh-Razavi & Kriegeskorte, 2014; Yamins et al., 2014; for review, see Cichy & Kaiser, 2019; Kietzmann, McClure, & Kriegeskorte, 2019). We extracted the activations of CNN layers for two purposes. First, we use them to identify the degree of selectivity of each object concept and how it increases from early to late layers. Second, we provide similarity matrices of the images as an additional resource for researchers. To this end, we used CorNet-S (Kubilius et al., 2018), a comparably shallow recurrent neural network architecture inspired by the ventral visual stream in the macaque brain. For each of the images contained in our database, we extracted the activations for all five layers in CorNet-S and converted them to vectors. For each layer, we then created a similarity matrix by computing the Pearson correlation coefficient between all pairs of vectors.

## Results

### 1,854 object concepts and 27 core high-level categories

The final list of concepts comprised 1,854 concrete objects, mass items (e.g. “sand,” “coal,” “gravel”) or other “things” (e.g. “footprint,” “fingerprint”), and objects spanned a wide range of different concepts. All object concepts are provided with their WordNet synset IDs, a link to an example image, concreteness ratings (Brysbaert et al., 2014) word frequency from several corpora (Brysbaert et al., 2012; Davies, 2008), category membership determined bottom-up through ratings and top-down through the WordNet hierarchy, definitions from WordNet, and others (for a full list, see S1 Table).

Based on the bottom-up categorization provided by human raters, 926 of these objects were rated as belonging to the 27 most common categories with at least 15 members (Table 1), with very little overlap between those categories (34 objects belonging to two categories). For simplicity, in the following we will refer to these 27 categories as the “core categories”. The distribution of category membership followed a rapidly decaying function, i.e. many of the remaining objects were categorized as belonging to small and very specific categories. While several of the 27 core categories provided by human raters overlap with those in classical category descriptions categories (Battig & Montague, 1969; Van Overschelde et al., 2004) and in WordNet (Fellbaum, 1998), other categories were unique to the ratings (“part of car,” “office supply,” “clothing accessory,” and “medical equipment”). A similar trend was observed for the top-down categorization provided by WordNet: 960 object concepts belonged to 23 of the 27 categories that were available in WordNet (the 24th category “car part” did not contain any subordinate entries). Of those, 118 objects belonged to multiple categories, which was mostly explained by the overlap introduced by the presence of subcategories (e.g. both insects and birds are animals).

**Table 1.** High-level object categories based on object concepts in THINGS database.

| Category | Synset | Frequency<br>(Mturk, bottom-up) | Frequency<br>(WordNet, top-down) | Overlap |
| --- | --- | --- | --- | --- |
| food | food.n.01 | 137 | 151 | 85 |
| animal | animal.n.01 | 95 | 174 | 94 |
| clothing | clothing.n.01 | 60 | 93 | 55 |
| tool | tool.n.01 | 51 | 50 | 25 |
| sports equipment | sports_equipment.n.01 | 46 | 22 | 15 |
| vegetable | vegetable.n.01 | 38 | 27 | 23 |
| vehicle | vehicle.n.01 | 37 | 66 | 28 |
| musical instrument | musical_instrument.n.01 | 33 | 32 | 30 |
| fruit | fruit.n.01 | 32 | 49 | 31 |
| body part | body_part.n.01 | 31 | 31 | 29 |
| dessert | dessert.n.01 | 29 | 11 | 9 |
| toy | toy.n.01 | 29 | 24 | 17 |
| container | container.n.01 | 28 | 139 | 23 |
| part of car | car_part.n.01 | 28 | 0 | 0 |
| weapon | weapon.n.01 | 28 | 20 | 17 |
| bird | bird.n.01 | 27 | 27 | 26 |
| furniture | furniture.n.01 | 27 | 32 | 18 |
| kitchen tool | kitchen_utensil.n.01 | 27 | 17 | 7 |
| office supply |  | 26 | n/a | 0 |
| clothing accessory |  | 21 | n/a | 0 |
| kitchen appliance | kitchen_appliance.n.01 | 21 | 10 | 8 |
| plant | plant.n.02 | 21 | 37 | 17 |
| insect | insect.n.01 | 20 | 16 | 16 |
| home décor | decoration.n.01 | 19 | 30 | 3 |
| medical equipment |  | 18 | n/a | 0 |
| electronic device | electronic_device.n.01 | 16 | 5 | 1 |
| drink | beverage.n.01 | 15 | 16 | 12 |

The 27 most common high-level categories as determined by Amazon Mechanical Turk workers providing categories for 1,854 objects and listing any category names more than 15 times. For the same categories, we determined the frequency for WordNet synsets and their subordinate members (hyponyms).

Since the dichotomies “animate – inanimate” and “man-made – natural” are frequently used high-level categorical distinctions, we counted the number of nameable objects that belonged to either of those categories. To our surprise, only around 11 % of the objects were animate – even when including humans and human body parts – while 89 % were inanimate. Similarly, for the dimension of naturalness – even when including processed food – only around 33 % of objects were natural, whereas 67 % were artificial. This demonstrates that, at least based on everyday object naming, a majority of object concepts indeed refers to artificial and inanimate objects. This result makes sense, given that humans are constantly surrounded by a large variety of inanimate and artificial objects whose distinction for everyday use is relevant. However, animals belong to one of the largest categories, are a rather homogenous class (e.g. most animals have faces) and may be more significant from an evolutionary point of view, which may contribute to their increased salience. Note that the significance of distinguishing between concepts by their linguistic use need not imply that categories with a more diverse set of concepts are more common; rather, it may prove to be useful to distinguish many different types of the same category if they have distinct functional significance.

A notable feature of the database is that a substantial proportion of objects (bottom-up categorization: 12.73 %, top-down categorization: 12.08 %) was categorized as belonging to one or several of the categories of edible items (“food,” “vegetable,” “fruit,” “dessert,” “drink”). Two more categories refer to kitchen items (“kitchen tool,” “kitchen appliance”), demonstrating the general importance of discriminating among food-related items.

### Relationship of core high-level object categories with semantic embedding

To determine the degree to which the high-level categories reflect their actual use in language, we visualized the similarity of concepts by running t-distributed stochastic neighborhood embedding (t-SNE, perplexity = 30, initialized with perplexity = 5, Maaten & Hinton, 2008) on the semantic embedding (i.e. synset vectors) and displaying the 27 core object categories in different colors (Fig 2A). The resulting visualization indicates that much of the categorization provided by humans is also mirrored in the synset vector similarity, although some structure is missed. To provide a quantitative basis for this result, we measured the selectivity of each concept by comparing the correlation of concepts within each of the 27 core categories to the correlation of those concepts with all other concepts (Fig 2B, all left bars). We repeated the same analysis for the high-level categories derived from WordNet (Fig 2B, all right bars). The results demonstrate a positive selectivity (i.e. correlation difference) for all 27 core categories, both for bottom-up and top-down categories (all *p* < 0.0054, Bonferroni-corrected over 27 categories, based on 5,000 Monte Carlo samples of categories). Overall, the selectivity of categories was higher for the “bottom-up categories” as compared to the categories defined based on WordNet (Δ*r*_*bottom-up*_ = 0.23, Δ*r*_*top-down*_ = 0.19, *p* < 0.0002).

**Fig 2.**
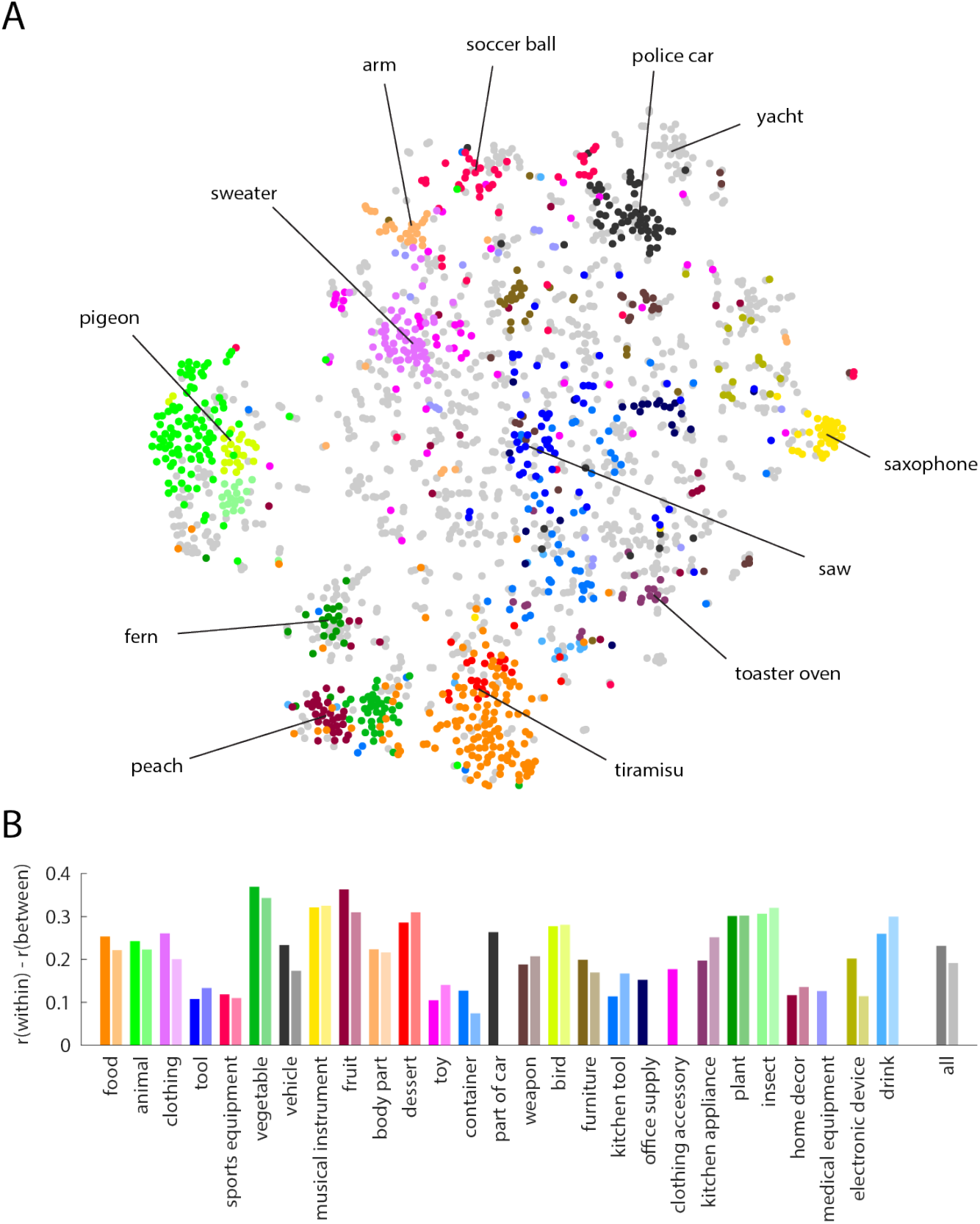
Category structure of THINGS object concepts. **(A)** Visualization of the semantic relationship of the 1,854 object concepts applying t-SNE to the semantic embedding, with the 27 core object categories depicted in different colors and example concepts highlighted. **(B)** Selectivity of the 27 core object categories, separately for bottom-up categorization based on responses of workers on Amazon Mechanical Turk (left bars, darker shades) and top-down categorization based on category membership in WordNet (right bars, lighter shades). Category selectivity was quantified by the difference in correlation of semantic embedding vectors of concepts within each category as compared to the correlation with concepts outside of the category (all *p* < 0.05, Bonferroni-corrected). Across all concepts, the selectivity for bottom-up categorization was higher than the selectivity for top-down categorization (*p* < 0.001).

### Object image database

For the object image database, we identified a total of 26,107 images (mean number of images per concept: 14.08). Of those images, 1,165 were selected from ImageNet. Example images for a small set of concepts are shown in Fig 3A. The mean image size was 996 × 996 pixels (< 1.8 % of images smaller than 500 pixels). The distribution of number of images per concept and pixel dimensions is shown in Fig 3B.

**Fig 3.**
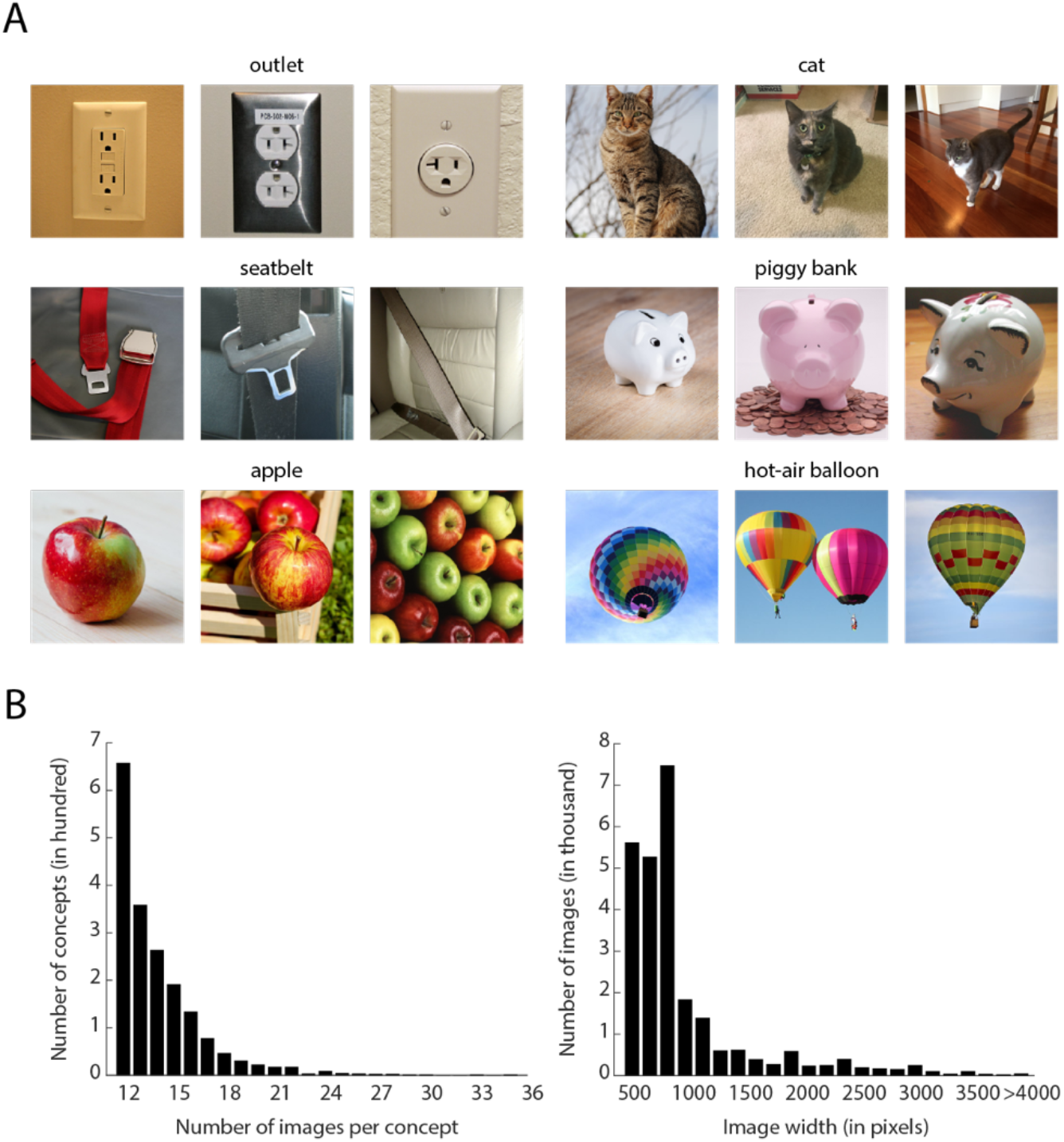
Examples and statistics of images in THINGS database. **(A)** Example images in THINGS database for different object concepts. **(B)** Histograms illustrating the number of images per concept (ranging from 12 to 35) and the image dimensions (peaking at 800 pixels).

To determine the degree to which individual object images are good members of a concept while still exhibiting visual variability, we fed all images through different layers of a deep convolutional neural network CorNet-S, as described in the *Methods* section (Kubilius et al., 2018). We predicted that at high layers, images within a concept would exhibit a larger similarity – as given by their Pearson correlation – than images between different concepts. In addition, we predicted this difference in similarity to be larger in the top layer (classification layer), but smaller in the bottom layer (V1 layer). Finally, for each of the 26,107 object images we investigated to what degree the activation of one object image could be used to predict other members of the same concept. The percentage of correct guesses is given by how many of the most similar images are populated by the *n-1* images of the same concept, i.e. excluding the reference image. We define the top-1 accuracy as the first *n-1* image ranks, the top-5 accuracy as 5 times that number, and the median rank as how many guesses are required for each other member of the concept. The results of these analyses are shown in Fig 4. For all layers, the correlation was higher within concept than between concept (classification layer: *r* = 0.51 within, *r* = 0.04 between; V1 layer: *r* = 0.39 within, *r* = 0.34 between, all *p* < 0.001, based on 1,000 randomizations) with the correlation difference increasing between the first layer and the classification layer (*p* < 0.001), demonstrating selectivity within each concept that increases across layers. The pairwise similarity was a good predictor of other members of the same concept for the classification layer (top-1 accuracy: 37.29 %, top-5 accuracy: 61.56 %, median rank: 29), but much less so for the V1 layer (top-1 accuracy: 1.25 %, top-5 accuracy: 3.51 %, median rank: 5,530), despite the network being trained only on a subset of concepts (334/1000 concepts overlap with THINGS, 212/1000 are subordinate examples of the concepts used in THINGS). Together, these results demonstrate that the object image database constitutes both a good representation of individual object concepts, while still exhibiting notable variation in low-level image properties.

**Fig 4.**
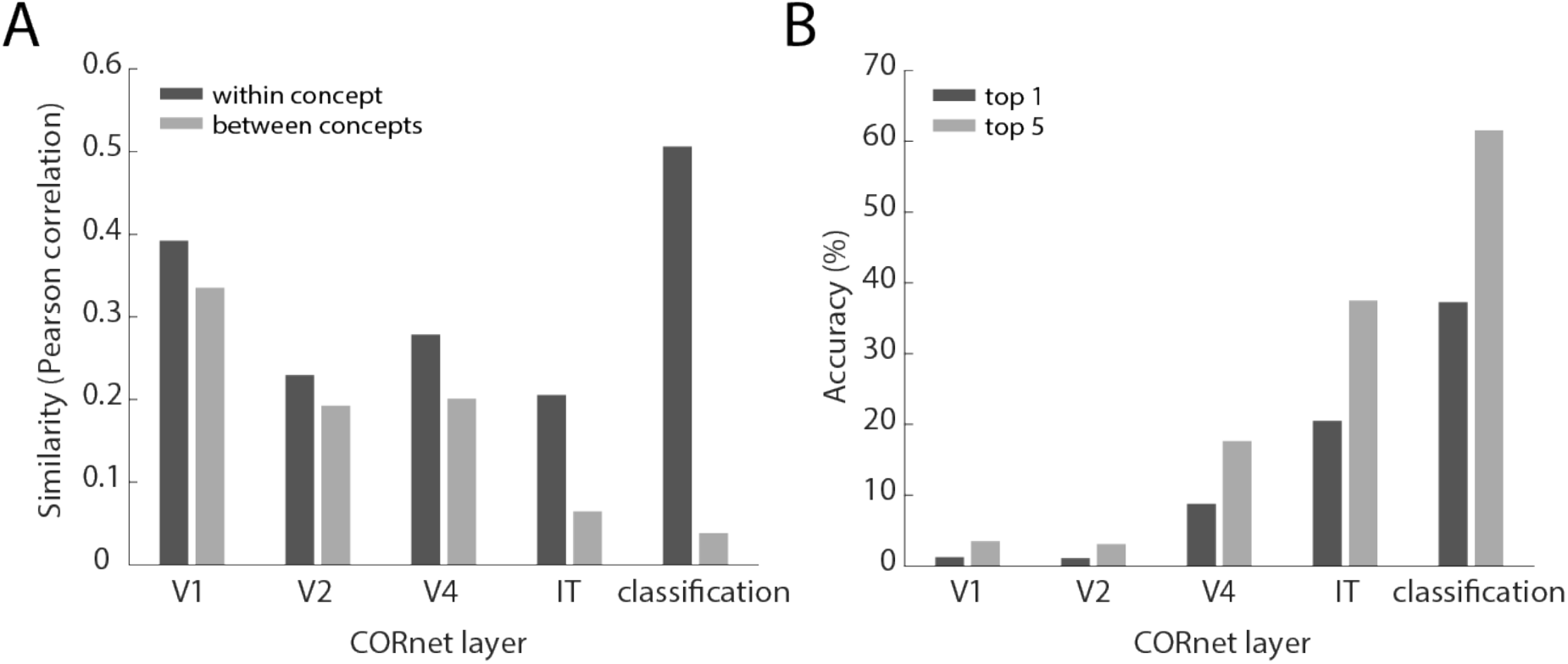
Selectivity of object images for different concepts. **(A)** Similarity of image activation vectors across different layers of the deep convolutional neural network CORnet (Kubilius et al., 2018). The first four layers of CORnet are named after brain regions in the macaque monkey. Similarity was estimated by the mean Pearson correlation of activation patterns in a given layer. “Within concept” refers to the similarity of all pairs of activation vectors of images of the same concept, while similarity “between concept” refers to the similarity of those images and all other images in the database. Higher selectivity is indicated by a larger difference in “within concept” and “between concept” similarities. **(B)** Predictive accuracy of activation vectors based on their pairwise Pearson similarity. “Top-1 accuracy” refers to the percentage of the most similar images that belong to the same concept as a reference image and is averaged across all accuracies for all images. “Top-5 accuracy” allows five guesses. Note that this procedure is not directly comparable to that used in typical machine learning applications (see main text for details).

## Discussion

Here we present a large-scale database of 1,854 diverse object concepts and 26,107 high-quality naturalistic images of those objects. For the object concepts, we identified 27 high-level categories that capture around half of those concepts with little overlap. We validated the categories and the object images by relating them to representations in a semantic embedding and a deep convolutional neural network, suggesting that the categories are meaningfully related to their use in language and that the object images represent largely distinct categories while varying in basic visual properties.

### Possible applications of the object concept and object image database

The purpose of this database is to provide researchers in the fields of psychology, neuroscience, and computer science with a resource they can use to systematically select object concepts or object images for their research. The availability of a large-scale resource has the advantage of providing a more standardized approach for the selection of object concepts and object images. In addition, if adopted more widely, it offers an increased level of comparability between different studies.

There are a wide range of potential applications of such a database, and we will briefly discuss only a few. In the field of psychology, studies in object recognition, categorization and semantic memory can use this database to identify a more representative set of concepts and provide naturalistic example object images with them. This would allow researchers to study the large-scale structure and format of mental representations of objects (Jozwik, Kriegeskorte, Storrs, & Mur, 2017; Konkle & Oliva, 2011; Long, Konkle, Cohen, & Alvarez, 2016; Zheng, Pereira, Baker, & Hebart, 2019), to reveal the degree to which categorization behavior (e.g. as measured in Kiran & Thompson, 2003; Kirchner & Thorpe, 2006; Rajalingham, Schmidt, & DiCarlo, 2015) generalizes to less commonly-used categories and more natural images, and to determine factors affecting the recall and recognition of concepts and images (Brady, Konkle, Alvarez, & Oliva, 2008; Klein, Addis, & Kahana, 2005; Konkle, Brady, Alvarez, & Oliva, 2010; Rotello, Macmillan, & Van Tassel, 2000). In the field of neuroscience, the object image database can form the basis for a more systematic large-scale assessment of object representations based on intensive measurements within a small number of subjects (Chang et al., 2018; Huth, Nishimoto, Vu, & Gallant, 2012; Kay, Naselaris, Prenger, & Gallant, 2008; Naselaris, Prenger, Kay, Oliver, & Gallant, 2009) and thus offer the basis for a dataset that is more representative than those commonly used as benchmarks in methods development and object recognition research (e.g. Kriegeskorte et al., 2008). In the field of computer vision, while the number of object images provided in this dataset is too small for training current deep convolutional neural networks, they can be used to test how wide a range of concepts are spanned by a given computational model. More generally, the concepts identified in this research could form a comprehensive set of labels for object classification at a level that is more comparable to humans (see also Mehrer et al., 2017).

### Comparison to previous work and existing databases

#### Selection of object concepts and object categories in previous work

For identifying and selecting object concepts for their studies, researchers have been using a variety of strategies, often focusing on a small set of basic-level objects (Edelman, Grill-Spector, Kushnir, & Malach, 1998; Eger, Ashburner, Haynes, Dolan, & Rees, 2008; Haxby et al., 2001; Rajalingham et al., 2015; Rice, Watson, Hartley, & Andrews, 2014), specific superordinate categories and examples within those categories (e.g. animals, tools, or vehicles, Bracci & de Beeck, 2016; Connolly et al., 2012; Gerlach, 2007; Hung, Kreiman, Poggio, & DiCarlo, 2005; Liu, Agam, Madsen, & Kreiman, 2009; Tranel, Logan, Frank, & Damasio, 1997), using objects that vary along assumed representational dimensions (e.g. animacy, man-made and biological objects, manipulability, real-world size, Caramazza & Shelton, 1998; Konkle & Oliva, 2011; Warrington & Shallice, 1984), and other criteria (for reviews, see Grill-Spector & Weiner, 2014; Mahon & Caramazza, 2009; Martin, 2007; Murphy, 2004). While these selection criteria are useful for the specific hypotheses at hand, the selected concepts are often not general enough to be of use for other research questions, requiring researchers to repeat the identification and selection process for their own purposes. In addition, the definition of what constitutes a category varies between studies, which affects the comparability of results and the conclusions that are drawn. The database presented in this work constitutes a wide range of concepts that allows their systematic and reproducible selection. For example, if the selection of a wide range of 50 object concepts is desired, researchers could apply cluster analysis to the synset vectors provided with this dataset and select one concept from each cluster.

A wider range and more standardized set of concepts to choose from is provided by category norms (Battig & Montague, 1969; Van Overschelde et al., 2004) or object property norms (Devereux, Tyler, Geertzen, & Randall, 2014; McRae, Cree, Seidenberg, & McNorgan, 2005), which can be useful for improving comparability between studies. Despite their common use, it is important to note that the concepts and categories in those norms were selected mostly based on their use in previous studies. Thus, they may span a rather selective set of objects and higher-level categories. In addition, they contain concepts at a level of description that may not align with how the object is commonly named (e.g. “python” vs. “snake”), and for some categories may not conform to the way humans would naturally group their members (e.g. “four-footed animal”). For those reasons, depending on the goal of the study, a more systematic approach for concept selection and category definition may be desired. In the future, we hope to provide more systematic category norms and object property norms for the large set of object concepts in the THINGS database, which would combine the benefits of previous efforts with those of the present work.

As discussed above (see *Methods*), a highly systematic compilation of object concepts is provided through WordNet (Fellbaum, 1998), a lexical database that stores words and their hierarchical relationship according to their meaning as so-called synsets. This format is very valuable for identifying unique meaning of ambiguous words (e.g. “bat” as a nocturnal animal or as a club used in sports), for merging synonymous words (e.g. “sofa” and “couch”), and for identifying more high-level categories in general. Some researchers have used WordNet to circumvent selection biases by randomly sampling from a broader set of concepts. For example, the 1,000 synsets in the ImageNet Challenge (Russakovsky et al., 2015) have become a standard set of object concepts in computer vision research, and a large part of them were sampled at random from WordNet. However, WordNet also contains a large number of concepts that can only be identified by experts (e.g. “tobacco hornworm,” “trogon”) or that cannot be distinguished easily by just looking at pictures of them (e.g. “black pine” vs. “red pine”). In addition, it is not always clear what level of categorization is the most useful for a given object (e.g. “canine” vs. “dog” vs. “poodle”) and to what degree the WordNet hierarchy translates to the everyday use of concepts and categories (e.g. in WordNet a “hydrant” belongs to the category of “discharge pipes”). The list of object concepts presented in this work addresses this challenge by identifying the level of description that matches their use in object naming, while still providing synsets to tie them to unique word meanings and relate them to the WordNet hierarchy.

Finally, another approach for the selection of object concepts in category selection is to label objects based on their natural appearance in photographs or movies and use these labels in later research (Huth et al., 2012; Russell, Torralba, Murphy, & Freeman, 2008). While this approach avoids a “top-down” selection bias, sampling of categories may lead to a “bottom-up” bias, by mirroring the statistics of the concepts found in the source stimulus set, thereby potentially overestimating the significance of frequent concepts and underestimating the significance of rarer ones.

#### Selection of object images in previous work

The format of visual presentation of objects is important for the study of visual cognition, including visual object recognition, memory, categorization, and naming. To this end, the use of standardized line drawings of objects has been a dominant approach (Snodgrass & Vanderwart, 1980). However, researchers have started to increasingly rely on the use of naturalistic object stimuli from photographs, in order to more closely match the conditions of real-world perception (Einhäuser & König, 2010; Felsen & Dan, 2005). Those naturalistic stimuli have been ranging from images of isolated objects cropped from their natural background (e.g. Baldassi et al., 2013; Haxby et al., 2001; Kiani, Esteky, Mirpour, & Tanaka, 2007; Kriegeskorte et al., 2008) to object renderings placed on naturalistic scenes (e.g. Yamins et al., 2014), object images with a naturalistic background (e.g. Rust & DiCarlo, 2010; Thorpe, Fize, & Marlot, 1996), multiple objects in naturalistic scenes (e.g. Peelen, Fei-Fei, & Kastner, 2009; Torralba, Oliva, Castelhano, & Henderson, 2006), and objects appearing in dynamic movies (e.g. Huth et al., 2012).

In recent years, numerous standardized object image databases have been published for psychological and neuroscience research (for review, see Brodeur et al., 2010; Brodeur et al., 2014), which most commonly consist of naturalistic images of objects cropped from their natural background, such as the Bank of Standardized Stimuli (BOSS, Brodeur et al., 2010). This approach has been and still is very valuable for the study of visual cognition and memory. At the same time, most of these image databases contain only one or very few examples of a given object (e.g. “lamp”) or only a small number of object concepts, but not both many concepts and numerous examples. In addition, the naturalistic context in which objects appear is known to be important to object processing (Oliva & Torralba, 2007), and there is evidence that the use of cropped images may overemphasize the role of shape features in measured neural representations (Bracci et al., 2017; Bracci & de Beeck, 2016; Coggan et al., 2016; Proklova et al., 2017; Proklova et al., 2016). For those reasons, depending on the research question, it is important to also consider the use of a wider range of object images embedded in their naturalistic context.

In computer vision, several large-scale object image databases exist that provide up to thousands of examples of individual objects in a naturalistic context (Deng et al., 2009; Griffin et al., 2007; Krizhevsky & Hinton, 2009). However, the size or quality of a large portion of images in those databases is not sufficient for their widespread use in psychology and neuroscience experiments and requires researchers to carefully and manually select candidate images to be of sufficient quality (Chang et al., 2018), which even after selection still involves trade-offs with respect to the size, aspect ratio and naturalness of those images. The THINGS database offers a comprehensive set of high-quality object images that should be of sufficient size for most applications in psychology and neuroscience research. While the number of exemplars in the THINGS database is notably lower than that in computer vision databases, the images can serve as a test set for assessing the generality of existing computer vision algorithms. For example, it is promising that a convolutional neural network trained on only a subset of the concepts in the THINGS database can still yield reasonable classification performance even without retraining and when using a comparably simple method for classification.

### Limitations of the THINGS database

#### Limitations in the selection of concepts

Many steps were involved in the definition of the list of 1,854 concepts, requiring choices during each step of the selection process. We laid out the exact choices at each step in great detail and provide the list of excluded words alongside the final list of concepts, allowing researchers to choose different inclusion and exclusion criteria if they wish. By the nature of the task, some of our choices were subjective and can be debated, and others involved a trade-off between efficiency and completeness. For example, for the initial selection based on concreteness we defined a cutoff below which we did not select any concept, likely excluding a small number of picturable and nameable objects. Further, we excluded objects that necessarily depict text. This makes sense given the goals of the present database, but it could arguably have been interesting to include them, given the known special role of text characters in the human brain (Cohen et al., 2000). However, the list of excluded text words is marked and available as part of the database. As another example limitation, one might argue that the list should have focused exclusively on objects and excluded items defined from mass nouns (e.g. “sand,” “coal”) or other items (e.g. “fingerprint”). We chose to include these items, because they are nameable, concrete and refer to entities beyond texture or surface material. Their exclusion would also have led to the exclusion of drinks, which in this database turned out to be one of the most common categories. However, researchers may choose to remove those items for their own purposes.

While we selected object concepts based on whether they were named consistently, this choice was based mostly on one reference image only and a relatively small number of participants per concept. Additionally, it required choosing another arbitrary cutoff for which concept to include or to exclude. As mentioned above, the goal of this database was not to create a definite set of all nameable object concepts. We chose to be rather inclusive in this step, so that excluded concepts would be those that were named inconsistently by participants. Links to all reference images are available as part of this database. In the future, researchers may choose a similar approach with a wider range of reference images and more behavioral responses to identify a more general set of object concepts.

Another limitation of our approach is that ultimately object concepts were selected based on a list of nouns. While this approach is common, it may bias the selection of concepts (and categories) towards representations that have a wider linguistic variability (e.g. different food items), which need not be representative of their mental representation. In theory, there are alternative approaches that circumvent the use of language, for example the ability to discriminate between different randomly-selected objects. However, without the use of language, such approaches would be challenging to carry out in practice. We chose a more pragmatic approach based on WordNet synsets and object naming that goes beyond typical approaches, by using a much wider range of nouns as a basis for concept selection. In the future, it might be possible to determine a larger, more representative set of concepts that humans can still distinguish without reverting to linguistic criteria.

#### Limitations of the object images

The object image database comes with limitations, as well. Most of these limitations are related to the difficulty of finding good examples of object images. Even though we intended object images to be of high quality and contain natural background throughout the database, for some concepts it was difficult to impossible to find good examples of images embedded in a natural background, so trade-offs with image quality had to be made. Similarly, the choice not to focus exclusively on images with single examples of objects but allow several instances of the same object in an image is debatable. On the other hand, this choice may in fact reflect a more natural form of object context.

Object viewpoint and object background might not vary to a degree sufficient to test viewpoint-invariant and background-invariant object representations. At the same time, image representations in early layers of the deep convolutional neural network turned out to be quite unspecific for individual object concepts, while representations at higher layers were quite specific. This indicates that the level of image variation effectively controlled for much of low-level processing while still providing relatively high degrees of concept specificity.

Since almost all object images were originally in jpeg-format, we chose to use the same image format for the image database. However, cropping images to square made it necessary to save images again with jpeg-compression, which may have introduced an additional loss in image quality that affects the frequency spectrum of the images. Thus, for future databases the use of lossless formats (e.g. png-format) may prove beneficial.

Finally, one limitation of this object image database is that it likely contains copyrighted material, which limits their use to academic purposes under fair use regulations and limits publication of example images in scientific journals (the images in Fig 3 were chosen to be non-copyrighted examples in THINGS that come from the public domain).

### Future directions

Apart from potential improvements for the choice of object concepts and object images, there are a number of avenues for future developments as part of the THINGS database. First and foremost, additional high quality images may be selected, cropped, and added to the database, ideally focusing on non-copyrighted examples such as those provided in the Open Images database (Kuznetsova et al., 2018). Second, for researchers interested in using representative cropped examples, existing databases may be amended for that purpose (e.g. Brodeur et al., 2014). Third, in addition to the object categorization provided by participants and through WordNet, the existing database can be amended with expert categorization, potentially further improving the correspondence to semantic embeddings. Fourth, those categories could be used to generate a comprehensive set of typicality ratings of objects concepts. Finally, the concepts can form the basis for the creation of feature norms similar to existing ones (Devereux et al., 2014; McRae et al., 2005) or explicit ratings of object dimensions (e.g. real-world size, animacy, manipulability, etc.). Together, we hope that the THINGS database is widely adopted by the scientific communities in psychology, neuroscience, and computer science, thereby broadening the use systematic and large-scale naturalistic research and further advancing the communication between these fields of research.

## Acknowledgements

This work was supported by the Intramural Research Program of the National Institutes of Mental Health (ZIA-MH-002909) and a Feodor-Lynen fellowship of the Humboldt Foundation awarded to M.N.H.

